# Ecological diversity exceeds evolutionary diversity in model ecosystems

**DOI:** 10.1101/2022.03.24.485441

**Authors:** Ilan Rubin, Yaroslav Ispolatov, Michael Doebeli

## Abstract

Understanding community saturation is fundamental to ecological theory. While investigations of the diversity of evolutionary stable states (ESSs) are widespread, the diversity of communities that have yet to reach an evolutionary endpoint is poorly understood. We use Lotka-Volterra dynamics and trait-based competition to compare the diversity of randomly assembled communities to the diversity of the ESS. We show that, with a large enough founding diversity (whether assembled at once or through sequential invasions), the number of long-time surviving species exceeds that of the ESS. However, the excessive founding diversity required to assemble a saturated community increases rapidly with the dimension of phenotype space. Additionally, traits present in communities resulting from random assembly are more clustered in phenotype space compared to random, though still markedly less ordered than the ESS. By combining theories of random assembly and ESSs we bring a new viewpoint to both the saturation and random assembly literature.

## Introduction

The diversity of competitive communities has been a focus of ecological theory since MacArthur’s seminal work on species packing and limiting similarity of competing species in a continuous trait space [MacArthur, 1970; Macarthur & Levins, 1967; MacArthur, 1965]. One of the enduring concepts to come out of this work is the question of whether natural communities are saturated (a community which is globally uninvasible) [Mateo *et al*., 2017]. In the extensive species packing literature, the diversity of ecological communities are usually studied once a population reaches an evolutionary equilibrium, which naturally implies that the communities are saturated.

While it is undeniably important to understand the patterns of “niche-packed,” saturated equilibrium, these ESSs (evolutionary stable states) represent the endpoint of evolutionary dynamics [Doebeli, 2011; Edwards *et al*., 2018] or the assembly of communities based on an infinitely diverse initial population. It has become more evident in recent years that evolutionary theory largely focusing on equilibrium dynamics is often limiting. Natural populations likely exist in non-equilibrium states [Olszewski, 2011], whether because of transient ecological dynamics [Fukami & Nakajima, 2011; Hubbell & Foster, 1986; Olszewski, 2011; Scheffer & van Nes, 2006], continually changing abiotic conditions [Herron & Doebeli, 2011; Kremer & Klausmeier, 2017; Svanbäck *et al*., 2009], or simply because an evolutionary stable state has not yet been reached. Therefore, it is important to study patterns of diversity for newly assembled communities of various sizes in relation to the robust theory surrounding species-packed, saturated equilibrium.

Notably, while competitive exclusion and limiting similarity have been used as explanations for the “paradox of the plankton” and the existence of a continuous distribution of species in trait space [e.g., Lehman & Tilman, 2000; Roughgarden, 1976], it has been repeatably shown that these coexistence continua are structurally unstable [Hernández-García *et al*., 2009; Leimar *et al*., 2013; Szabó & Meszéna, 2006]. In all structurally stable formulations of the competitive exclusion model, communities with distinct species (as defined by their phenotype) are expected.

The importance of studying randomly assembled communities as a proxy for natural ecosystems was first appreciated by May [1972]. May was able to determine the stability of large communities when species interactions were defined by random interaction matrices [May, 1972]. Random matrix theory has continued to be productive in helping to understand the ecological stability [Allesina & Tang, 2012, 2015; Fowler, 2009; Lawlor, 1980] and feasibility (when there is a stable coexistence of all species in the community – i.e., no species in the community go extinct) of large communities [Barabás *et al*., 2016; Dougoud *et al*., 2018; Serván *et al*., 2018; Song & Saavedra, 2018]. However, for these types of models, by directly defining species based on a random interaction matrix, rather than allowing interactions to be an emergent property of trait-based competition, community saturation can only be defined in the context of the species present in the interaction matrix. When instead defining species interactions by trait-based competition, community saturation becomes a characteristic of the ecosystem as well as the species present. A phenotype-based competition function is not only more realistic than randomly generated interaction matrices as the phenotypic competition introduces mechanistically-justified correlations into the interaction matrix (as the first-level approximation beyond the completely random matrix), it also allows us to have a clear picture of the evolutionary dynamics and to compare ecologically assembled diversity to the evolutionary stable state (ESS) produced by such an evolution.

Here, we use classic Lotka-Volterra ecological dynamics with species interactions defined by competition in an *n*-dimensional trait space to investigate the feasibility and stable diversity emerging from randomly assembled communities. We compare the emergent diversity of these random communities to the niche-packed, saturated evolutionary equilibrium for that system. The main question we address is how diverse of a randomly assembled founding population is required to result in a stable community with at least the same diversity as the ESS. We follow up by comparing the emergent diversity from different types of assembly processes and investigating how efficiently randomly assembled communities divvy up the available niche space.

## Methods

### Ecological Dynamics

In order to answer these questions we use a model based on classic Lokta-Volerra ecological dynamics with a *d*-dimensional phenotype, **x**, to determine a species’ carrying capacity and competitive ability. For simplicity we set the intrinsic growth rate of each species, *r* = 1 and only consider symmetric competition (*α*(**x, y**) = *α*(**y, x**) ∀**x, y**). While these are both significant simplifications of the dynamics, they are reasonable, and oft-used, assumptions to facilitate an initial understanding of these dynamics.

The carrying capacity of each species is defined as 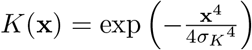 and species competition is modeled by a Gaussian competition function, 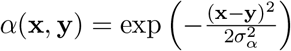. Therefore, *α*(**x, x**) = 1 and *α*(**x, y**)|_x≠y_ < 1. Together the ecological dynamics for species *i* in a population of *H* total species are as follows:

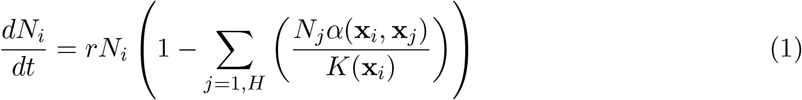

Here we chose to use a Gaussian competition function and a quartic carrying capacity function (*σ_α_* = 0.6, *σ_K_* = 1 unless otherwise noted). This system is a commonly used model to study diversification [Doebeli & Ispolatov, 2014; Doebeli *et al*., 2017; Rubin *et al*., 2021] and is a mathematically simple and structurally stable description of diversification dynamics. Using a higher order carrying capacity (and thus comparatively flat-shaped) function compared to the competition kernel both naturally restricts the viable phenotype space to an area around the origin (roughly between –2 and 2 for the parameters chosen) and guarantees the presence of a branching point at the origin [Baptestini *et al*., 2009; Doebeli, 2011], regardless of the choice of *σ_α_*.

The ecological dynamics can be rewritten without the explicit phenotypes as

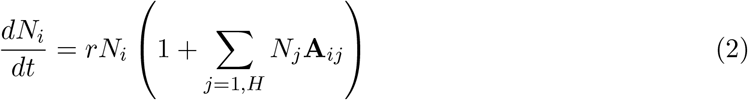

where the interaction matrix **A***_ij_* = –*α*(**x**_*i*_, **x***_j_*)/*K*(**x***_i_*). This equation is the classical formulation of multi-species Lotka-Volterra dynamics where the interactive nature of the species are defined directly, rather than with explicit phenotypes. In a competitive community, like the systems we will study here, all elements of the interaction matrix, *A*, are negative. Including positive elements in a interaction matrix can be instead used to model more complicated systems including mutualistic or predatory relationships between species.

Of particular note, the symmetric competition function, *α*(**x, y**), we are using here does not mean that the resulting interaction matrix, *A*, need be symmetric and in practice it rarely is. However, as our competition function is indeed symmetric, there is a single globally stable equilibrium for any community [Hernández-García *et al*., 2009], which in general includes finite and zero populations. Therefore, for symmetric competition an equilibrium population is the only possible outcome and no complex ecological dynamics (e.g., periodic or chaotic changes in population size) occur.

### Ecological simulations

Communities were assembled by generating *H*_0_ species, each with a randomly chosen phenotype **x** ∈ (–2,2)^*d*^. We were then able to solve for equilibrium population of the community by **N**\* = –*A*^-1^ * **r** (here, for simplicity we assume **r** = 1) and thus determine it is a feasible population if 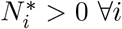. As mentioned above, the feasible equilibrium is always globally stable, so any feasible equilibrium represents the endpoint of the ecological dynamics.

However, if the randomly assembled community is not feasible, we are unable to analytically solve for the make-up of the final, stable population. Thus, we solved for the final stable community by numerically integrating the system of ODEs using the Runge-Kutta-Fehlberg method. Each simulation was integrated until *t* = 10^12^. Any species with *N_i_* < 10^-8^ was deemed extinct. The feasibility of the final simulated community was also calculated to determine if the ODE was run for enough time to reach a stable configuration.

### Calculation of the ESS

In order to compare communities to the Evolutionary Stable State (ESS) for the given environment, evolutionary dynamics of each system were also calculated. Evolutionary simulations were accomplished using an adaptive dynamics framework. As the evolutionary dynamics are not a focus of this paper, and we are using a standard model and technique, the evolutionary dynamics will not be explained in further detail. Please see Doebeli [2011]; Rubin *et al*. [2021] or the source code for these simulations included in the supplementary materials for information on the adaptive dynamics of this system. Notably, while we used adaptive dynamics as we felt it was the fastest and most accurate way to determine the ESS of each system, a successive invasion simulation like the one used by D’Andrea *et al*. [2019]; Rael *et al*. [2018]; Sasaki & Ellner [1995] would deliver exactly the same ESS results.

## Results

### ESS and Evolutionary dynamics

In order to find the saturation of point of each system we simulated the ESS using adaptive dynamics. As mentioned above, because the carrying capacity function is of a higher order (i.e., flatter) than the competition kernel, there is always a branching point at the origin, regardless of the choice of *σ_α_*. However, the presence of a branching point is not indicative of the diversity of the final populations. Using adaptive dynamics, the ESS for was calculated for varying widths of the Gaussian competition kernel from *σ_α_* ∈ [0.35, 1.5]. The ESS contains 2 species for any *σ_α_* ≳ 0.691, 3 species for 0.691 ≳ *σ_α_* ≳ 0.525, 4 species for 0.525 ≳ *σ_α_* ≳ 0.460, and an increasing diversity from there as *σ_α_* decreases.

With competition in dimensions > 1, the ESS has the same values as the equivalent 1-dimensional system, with the species laid out approximately in an *n*-dimensional lattice hyper-cube. This results in the diversity of the ESS in n-dimensions equal to the ESS of the same parameterization in 1-dimension to the raised to the dimension. For the rest of the discussion we will focus on communities based on *σ_α_* = 0.6 corresponding to an ESS with diversity *H_ESS_* = 3*^d^* [e.g., Figs. 1, C1].

**Figure 1:**
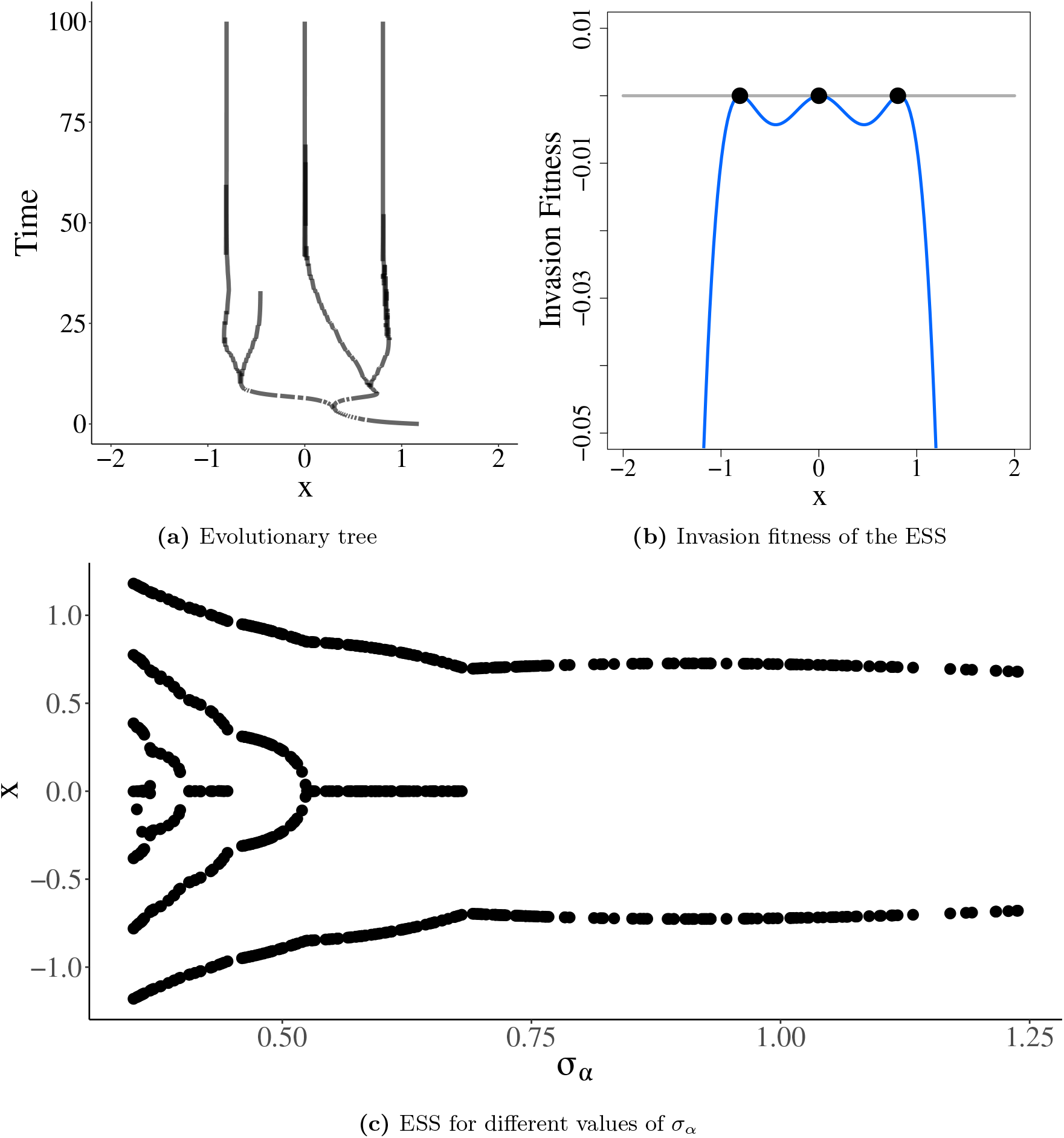
Evolutionary Stable States. Evolutionary stable states were calculated using adaptive dynamics. Panel A shows the evolutionary dynamics over time and panel B the invasion fitness and ESS both for *σ_α_* = 0.6. Panel C is a diversification diagram showing the simulated ESSs for randomly chosen *σ_α_* ∈ [0.35, 1.25]. Adaptive dynamics simulations were run for 10^7^ time steps, including 10^5^ branching mutations. The species were then clustered using a K-means clustering algorithm with a minimum distance equal to 0.1. As noted in the text, these ESS are not particular to adaptive dynamics but could also have been generated using a algorithm of ecological dynamics including successive invasions of random species. ESSs in higher dimensions are comprised of the same trait values in 1 dimension laid out in an n-dimensional lattice. For a representation of the ESSs for *σ_α_* = 0.6 in 2 and 3-dimensions see Figure C1.

**Figure 2:**
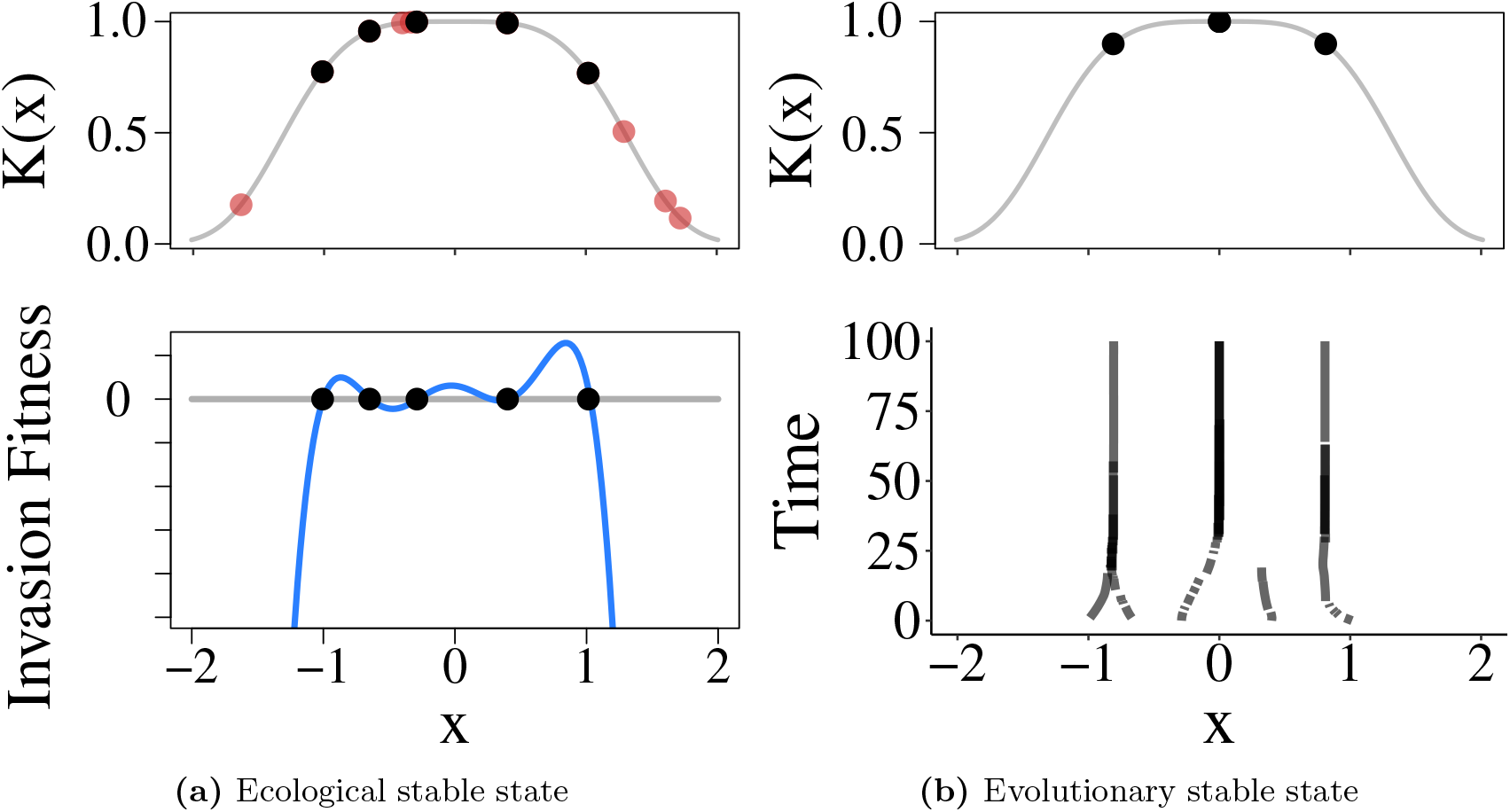
Example of 1D ecological and evolutionary stable states. An example of ecological and evolutionary stable states for a randomly assembled population with one dimensional phenotype (*x*). Founding population contains 11 randomly chosen species. Panel A shows the population after 10^12^ time steps and their carrying capacities. Species that survived are shown in black and those that went extinct in red. The blue curve shows the invasion fitness (per capita growth rate of a small mutant) of the ecological stable community. Panel B shows the evolutionary dynamics and evolutionary stable state (ESS) with the same community after evolution (as simulated with adaptive dynamics) is run for 100 evolutionary time steps.

### ESS is the endpoint of evolution

As mentioned above, the ESS represents the potential endpoint of the community on evolutionary timescales. Comparing the equilibrium diversity of communities on ecological and evolutionary timescales can be insightful into natural populations that do not strictly exist in equilibrium. As nicely summarized by Edwards *et al*. [2018], communities not at an ESS will experience selective pressure towards the ESS [however, see Doebeli & Ispolatov, 2014; Doebeli *et al*., 2017; Rubin *et al*., 2021, for discussions of non-equilibrium evolutionary dynamics]. In evolutionary time, super-saturated communities will likely experience the extinction of certain species and collapse to the ESS [e.g., 1]. Similarly, under-saturated communities represent ecological opportunity in the ecosystem which drives adaptive diversification toward the ESS [Schluter, 2000].

However, solely focusing on ESS minimizes the fact that evolution is ongoing. Studying ecological assembly processes has been a fundamental part of ecological theory [Serván & Allesina, 2021], both to understand the founding of nascent communities as well as an analogy to constantly changing abiotic conditions and the opening of new ecological opportunities. Comparing the diversity emergent from a random assembly process to the diversity of the ESS (*H_ESS_*), allows us to study the saturation of ecological communities before they reach an ESS.

### Randomly assembled saturated communities are rarely feasible

When communities are made simply from a random assortment of competitive species (each species has a randomly chosen phenotype and competes in an *n*-dimensional trait space), saturated (*H* = *H_ESS_*) or super-saturated (*H* > *H_ESS_*) communities are almost never feasible [Fig. C6]. At least one of the species will almost always go extinct. This follows simply from the theory of limiting similarity as there is a limited range of viable phenotypes. However, considering just the feasibility of random communities is fairly limiting as it does not allow for the natural assembly of stable communities through species invasion and extinction [Levine *et al*., 2017]. Thus, in order to determine the stable diversity (*H*\*) that results from a randomly assembled founding population, the ecological dynamics must be simulated.

For a longer discussion of the feasibility of randomly chosen species, including the effect of the dimension of the phenotype space on the probability of feasibility, please see Appendix A.

### Maximal ecological diversity

We first consider the canonical case of “top-down” community assembly [Serván *et al*., 2018; Song & Saavedra, 2018], where *N* randomly selected species are placed together at a small population size and allowed to equiliberate to a stable community via numerical simulations of the Lotka-Volterra ecological dynamics.

As expected, as the diversity of the initial founding population (*H*_0_) increases, so does the expected diversity of the final community (*H*\*) [Figure 3]. However, this increase in expected feasible diversity is also clearly a sub-linear function, such that subsequent increases in founding diversity lead to a marginal increase in the diversity of the final community. While we make no mechanistic hypotheses to the specific shape of this diversity expectation curve, there is a visually proficient fit to a Michaelis-Menten regression [Figure 3]. This is suggestive that the final diversity of a community is asymptotic as a function of the initial founding diversity, and therefore there being a maximal diversity possible for each system when communities are built through random assembly.

**Figure 3:**
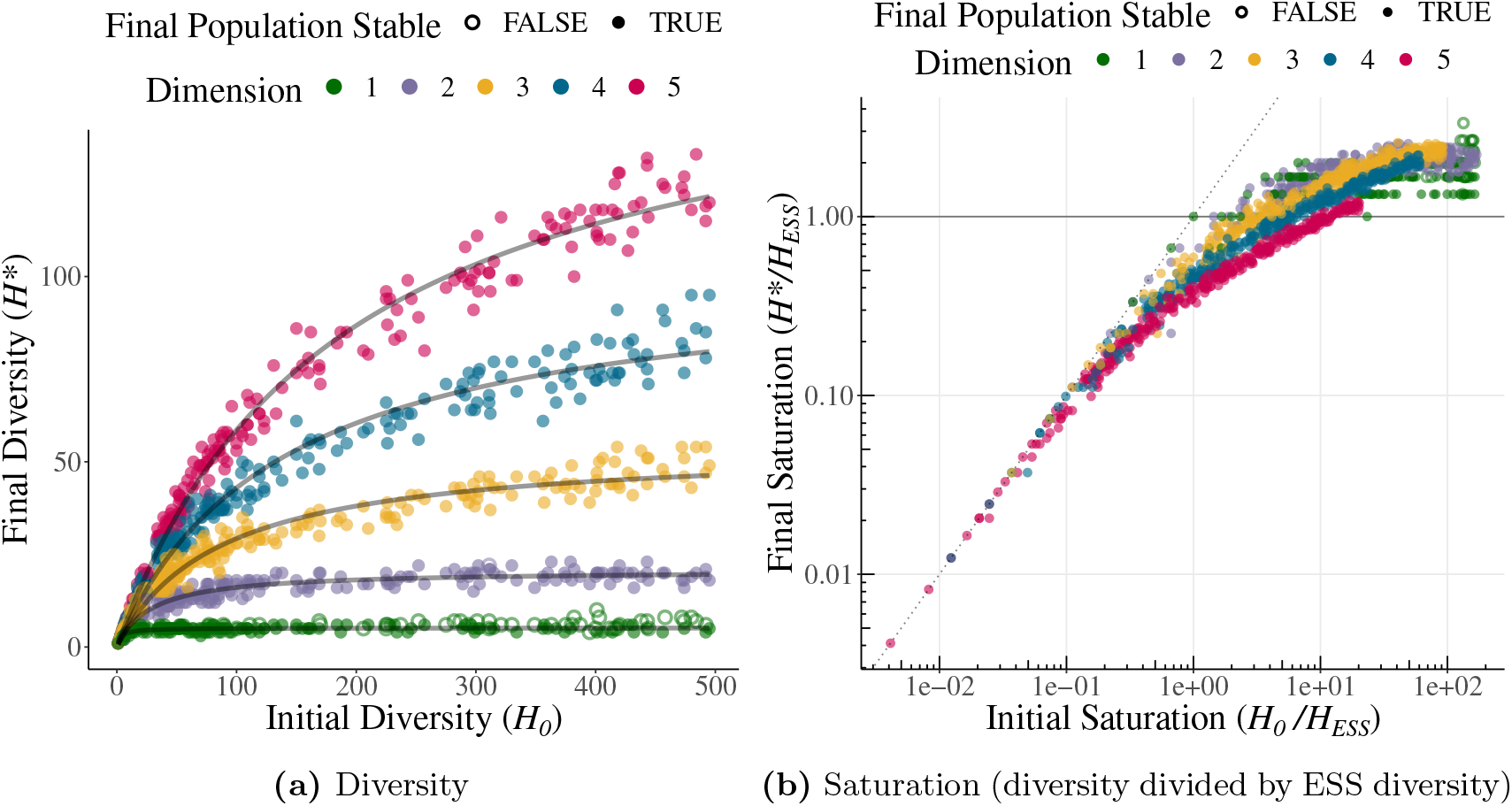
The final stable diversity and saturation of a randomly assembled community. Communities were assembled with random species in 1-5 dimensions. Ecological dynamics were run for 10^12^ time steps. Hollow points represent communities that did not stabilize to a feasible community by the end of the simulation and still contain at least one species that will eventually go extinct. Panel A shows the final diversity of communities as a function of their founding diversity. Grey lines are a regression using a nonlinear least squares fit to Michaelis-Menten form 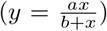. Regressions were fit only to communities that have stabilized by the end of the simulation. The Michaelis-Menten curves are not mechanistic, and were not chosen to model the ecological dynamics in any way, but are asymptotic curves (asymptote equal to regression parameter *a*) that are a good visual fit for the data. Panel B shows the saturation of the same communities as a function of the saturation of the founding community. Saturation is measured as the diversity divided by the diversity of the ESS (*H_ESS_* = 3*^d^*). Only communities with founding diversity of up to 500 species are displayed for panel A. For panel B communities were built with up to 500 species for dimension 1, 1500 for dimension 2, 2500 for dimension 3, and 5000 for dimensions 4 and 5.

In one dimension the maximally feasible diversity is twice the ESS. This occurs when each species present in the ESS is replaced by two species, balanced on either side of the ESS species in phenotype space. The invasion of any third species with a similar phenotype will cause either one or both of those coexisting species to go extinct. While this can be confirmed analytically for a 1 species ESS, for higher diversity states it is confirmed by extensive numerical simulations.

In higher dimensions more complex, and dense, patterns of species in phenotype space are possible. It is not hard to manufacture very saturated communities comprised of species laid out in symmetric patterns (like hyper-spheres and hyper-cubes) surrounding each ESS point in phenotype space. The feasibility of these synthetic communities are all dependant on their symmetry in phenotype space and are extremely sensitive to small perturbations of the phenotypes present.

Notably, while our simulations resulted in an ever increasing final community diversity with increasing founding diversity, very large founding populations will likely equiliberate to the ESS, even without considering evolution. This is a scenario that was been studied often [Sasaki & Ellner, 1995; Scheffer & van Nes, 2006] and is another method of numerically determining the ESS. However, we only saw high diversity assemblages result in the ESS for 1-dimensional simulations with *σ_α_* = 1.0 (in this case the *H_ESS_*=2) despite simulating communities with a diversity over two orders of magnitude larger than the ESS [Figure 3]. Thus, while the ESS is the high diversity limit of random assembly, such diverse assembly processes that always result in the ESS are unlikely to play a role in the formation of natural communities.

### Community saturation

To understand the pattern of the diversity of community assembly it is also important to consider the saturation (defined here as *H/H_ESS_*) of both the initial and final communities in addition to just the diversity of these communities. In these randomly assembled communities, community saturation seems to asymptotically approach a maximal saturation of around two to three times the ESS, regardless of dimension or niche width (defined by *σ_α_*) [Figure 3]. This suggests that while increasingly complex and dense patterns of coexisting species are easy to manufacture in higher dimensions, these patterns are unlikely to arise, at least with any significant frequency, via random assembly.

While randomly assembled communities have approximately the same maximal saturation regardless of dimension, creating a saturated or super-saturated community is increasingly difficult as dimension increases [Figures 3, 4]. The median number of species needed to assemble a saturated community increases from 2.17 times the ESS for 1-dimension (*H*_*ESS,d*=1_=3, median *H*_0,*d*=1_=6.5 for saturated communities) to 10.7 times the ESS in 5-dimensional trait space (*H*_*ESS,d*=5_=243, median *H*_0,*d*=5_=2597 for saturated communities). This means that for 5-dimensional trait space 400 times the founding diversity is required to assemble a saturated community compared to communities competing in 1-dimensional trait space. As the diversity of the ESS continues to grows exponentially with higher dimensional trait space, the even greater founding diversity necessary in higher dimensions to assemble a saturated community becomes intractable very quickly.

**Figure 4:**
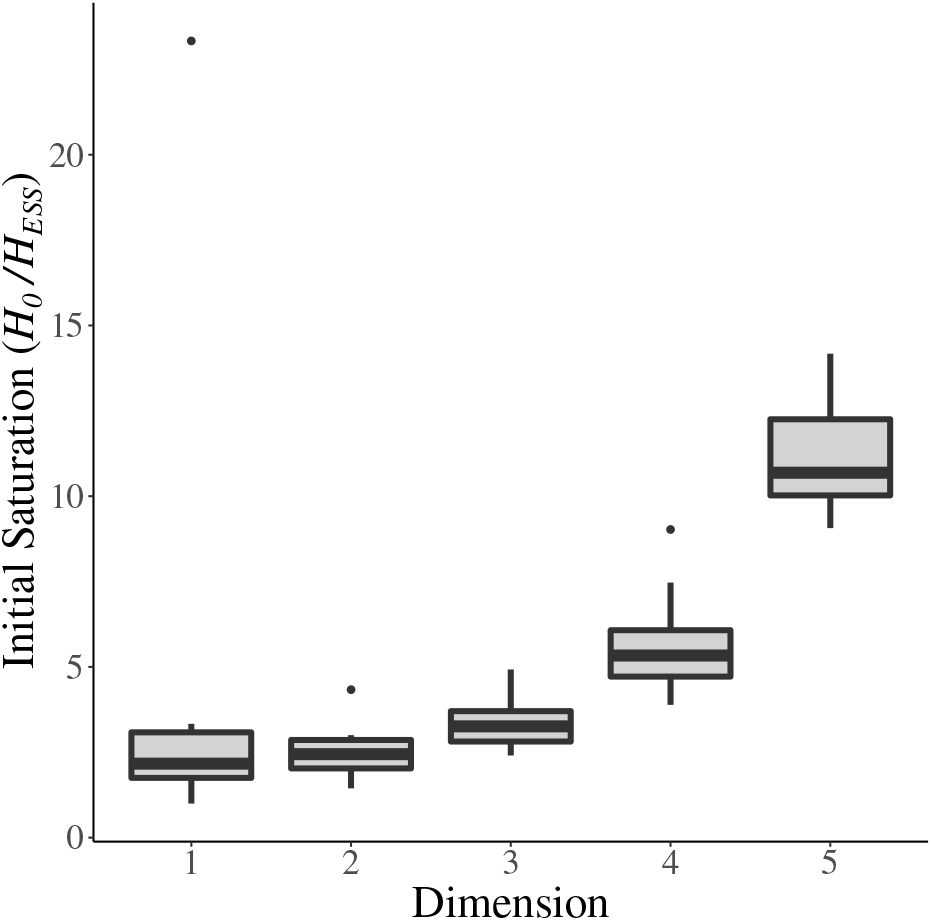
Initial saturation of a randomly assembled community necessary for the saturation of the final community. Here we show the saturation (diversity divided by the diversity of the ESS) of the initial, randomly assembled communities that end up stabilizing at approximately (95% to 105% of saturation) the saturation point for the given system. To create a saturated community from a random assembly of species, a more saturated initial founding population of species is required as dimension increases. The boxplots show the median, first and third quartiles, as well as extreme values.

### Diversity of top-down vs. bottom-up community assembly

In addition to the top-down community assembly we describe above, we also consider community assembly from a bottom-up process. In a bottom-up assembly processes random species are added to the community sequentially, instead of placing all *N* species in the system at once (top-down assembly) [Coyte *et al*., 2021; Diaz-Colunga *et al*., 2022; Serván & Allesina, 2021]. The community is initiated with two random species. Ecology is simulated for 10^10^ time steps or until the community stabilizes (|*dN_i_/dt*|< 9 × 10^-9^ ∀*i*) at which point another randomly chosen species is added. The simulation ends when *N* species (including the initial 2) have been added to the community.

Communities assembled from the bottom-up displayed roughly the same diversity expectations as those assembled from the top-down, except randomly-assembled bottom-up communities tend to be slightly less diverse than communities assembled from the top-down with the same number of species [Figure 5].

**Figure 5:**
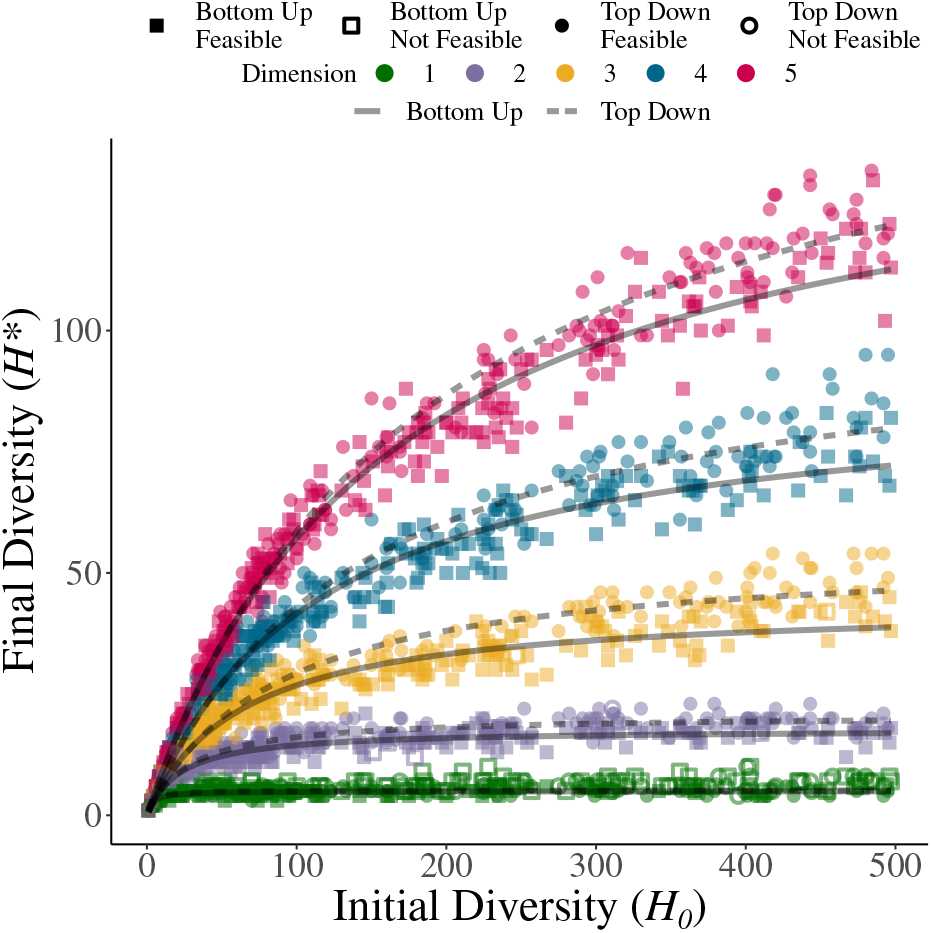
Top-down versus bottom-up community assembly. Top-down communities were assembled by selecting N random species and initiating ecological dynamics with all species together. Bottom-up communities were assembled sequentially one species at a time until N species were allowed to invade the community. The ESS in one dimension is made up of 3 species and increases exponentially with dimension (*H_ESS_* = 3*^d^*). The grey lines are a regression using a nonlinear least squares fit to Michaelis-Menten form. Regressions were fit only to communities that have stabilized by the end of the simulation. The solid lines and squares represent bottom-up communities; dashed lines and circles are top-down.

Serván & Allesina [2021] show that in competitive communities the community generated via a top-down assembly process can also always be reached sequentially (bottom-up) if drawing from the same species pool. However, multiple final, stable communities are possible with the same species pool. The priority effects and historical contingencies [Chase, 2003; Fukami & Nakajima, 2011] emergent from the specific order of invading species in a bottom-up assembly often results in an alternative stable community compared to one built top-down.

Here, the increased diversity in top down communities is likely because of complex patterns of species in phenotype space that require a specific order of species invasions to form sequentially [Sigmuiud, 1995]. These are configurations of species in which at least some of the pairwise combinations of species are unable to coexist in isolation but are stabilized by the presence of another species in the community. These multi-species consortia either have to be built up in a specific order of sequential species or can only exist when all species are present at the same time (e.g., via top-down assembly). Thus, despite the higher diversity community being accessible via a sequential assembly processes, some of the time bottom-up assembled communities get stuck in lower diversity stable communities.

However, as noted above, the diversity difference between the two assembly processes is fairly minor. Initial diversity and the dimension of competition are much stronger indicators of both final community diversity and saturation than the nature of the assembly processes. While we did not test other, more complicated assembly processes comprising of invasions with more than one species at a time [Diaz-Colunga *et al*., 2022; Gilpin, 1994], it logically follows that the final diversity of these communities will be somewhere in between those built in a purely top-down or bottom-up fashion.

### Trait dispersion of randomly assembled communities

A key feature in species-packed equilibria are evenly spaced species in niche space to efficiently divvy up the available niche space [Macarthur & Levins, 1967; Meszéna *et al*., 2006]. To investigate this we calculated the pairwise distances (in phenotype space) of each species to every other in the community. The “dispersion” of the community was then measured by the coefficient of variation (standard deviation divided by the mean) of the distance of each species to its nearest neighbor (*CV_NN_* = *σ/μ*) [Barabás & D’Andrea, 2016]. As the ESSs for these systems take the form approximately of an *n*-dimensional lattice (the distances between rows/columns of the lattice are not necessarily equal when the *H_ESS_* ≥ 4*^d^*, with species near the origin being slightly closer together), these ESSs are very under-dispersed (more regularly spaced in trait space than would be expected at random). The coefficient of variation of the distance to the nearest neighbor separation in these lattice-like ESSs is equal to (in systems with an *H_ESS_* ≤ 3*^d^*) or close to (in communities with an *H_ESS_* ≥ 4*^d^*) 0.

While randomly assembled communities were more dispersed in trait space than the lattice-like configuration of the ESS, they were also distinctly less dispersed than random configurations [Figure 6]. This was further confirmed in one dimension by using Welch’s t-test to statistically assess the dispersion of both top-down and bottom-up communities versus random traits of the same diversity [Table B1]. For both community assembly processes (top-down and bottom-up) and all levels of feasible diversity (3-6 species), the randomly assembled communities have significantly under dispersed traits by this measure compared to both uniformly and Gaussian distributed traits (*p* < 0.05). This means that even without evolution, communities will self-organize to efficiently utilize the available niche space solely through the extinction of overlapping species.

**Figure 6:**
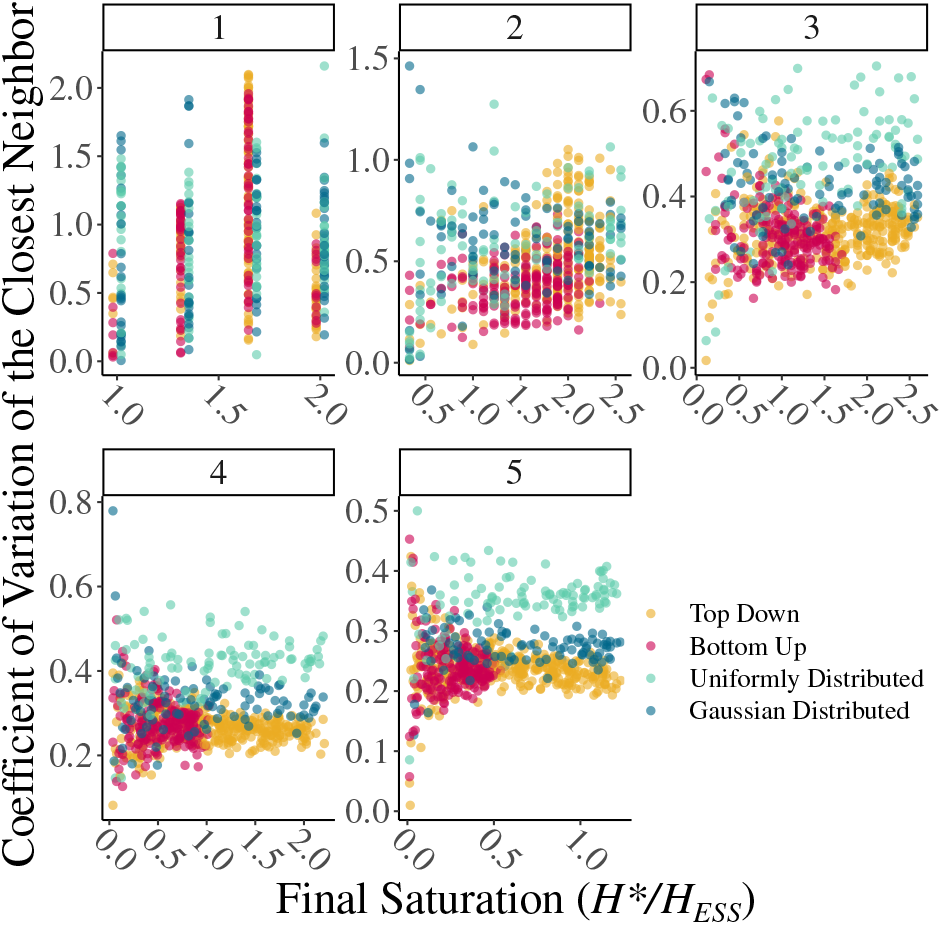
Evenness of randomly assembled communities in niche space. The pattern of trait dispersion was measured for randomly assembled communities using the coefficient of variation of the distance to the nearest neighbor (*CV_NN_*) for each species in the community. A value of *CV_NN_* = 0 corresponds to a perfectly ordered distribution of species, or a lattice in trait space. A very high *CV_NN_* would correspond to a very disordered community that has little organized structure. Colors represent the method of community assembly with yellow and red representing top-down and bottom-up randomly assembled communities; green and blue are the null expectation of uniformly (∈ [–2, 2]) and Gaussian distributed (*μ* = 0, *σ* = 1) traits. Bottom-up communities were built with up to 500 random species. Top-down communities were built up to 500 species for dimension 1, 1500 for dimension 2, 2500 for dimension 3, and 5000 for dimension 4 and 5. Ecological dynamics run for 10^12^ time steps. Only communities that stabilized to a feasible community when the simulation was terminated are shown. Results for 1-5 dimensional trait spaces are shown in each panel.

Patterns of trait dispersion become especially clear as community saturation increases. As saturation increases, species become more and more evenly spaced in trait space. Interestingly, at least for the sizes of communities we were able to simulate, the *CV_NN_* seems to approach ≈ 0.25, for all dimensions greater than one. This is somewhat surprising, as it is easy to assume that communities become increasingly over dispersed as they become more saturated. However, it seems that in higher dimensions, completely symmetrical and even distributions of species are unlikely to form.

This finding of common under dispersion of traits in randomly assembled communities mostly corroborates Barabás & D’Andrea [2016], who found that in simulated communities of 51 species, those that have evolved due to competition are significantly more ordered compared to the null expectation. While they were unable to find a statistical significance, they also state that randomly assembled communities without evolution seemed under-dispersed than the null expectation.

Also of note, top-down and bottom-up communities of the same diversity were not found to differ in this measure of trait spacing. This means that while bottom-up assembly leads to a slightly less diverse community on average, once established, communities of a certain diversity are indistinguishable based on assembly process.

### Transient extinction

We ran the ecological dynamics for an extremely long timescale (10^12^ time steps) which ensured that almost all simulations eventually settled to a stable community. However, even after this long time, some communities did not yet stabilize. It is a known phenomenon that extinction can be “exceedingly slow” [Hubbell & Foster, 1986]. Scheffer & van Nes [2006] found that extinction transients can cause the persistence of similar species causing “clumping” of species in trait space. We can confirm the same phenomena and thus unsurprisingly, very saturated communities, where there are significant numbers of very similar species, had the longest lasting transient extinction. For an example of the ecological dynamics of a single community with transient extinction please see Figure C5.

Not all cases of transient extinction was caused by the clumping of similar species. In rare cases nearly symmetric patterns of species in niche space, usually centered on a singular point in the ESS, resulted in a nearly “balanced” configuration and transient dynamics. however, these cases were rare and were not identified in many communities generated via random assembly.

## Discussion

One the central problem in ecology and evolutionary theory that has seen a resurgence in interest in recent years, is understanding the characteristics of an ecosystem that lead to the coexistence of many species. Previous work on the topic has largely focused on the distribution of species at an ESS [e.g., MacArthur, 1970], coexistence due to transient dynamics [Scheffer & van Nes, 2006], or community assembly through the creation of random interaction matrices in the Lotka-Volterra model framework [Allesina & Tang, 2015; Barabás *et al*., 2016]. Here we have shown that while the diversity of the ESS of a system is certainly important, communities built through random assembly can be both unsaturated or super-saturated compared to that ESS depending on the diversity of the random assembly. However, for every system as long as the founding community is diverse enough, the resulting community is all but guaranteed to be saturated. This is true for both top-down and bottom-up assembly processes (though communities created with a top-down processes are on average slightly more diverse than those create from the bottom up).

Perhaps most importantly, the founding diversity necessary to create a saturated community actually increases faster than the increase in the diversity of the ESS as dimension increases. This means in natural communities that are likely competing in high dimensional phenotype space [Barbier *et al*., 2021; Doebeli *et al*., 2017; Débarre *et al*., 2014; Ingram *et al*., 2018; Pacala & Roughgarden, 1982], the generation of a saturated community through random assembly is essentially impossible. In higher dimension, evolution and frequency dependant selection are required to build saturated communities that efficiently divide trait space into regular niches. Even with very large founding diversity, communities built through random assembly retain some of the structural inefficiencies in trait dispersion from the complex and random assortment of species in comparison to the lattice-like ESSs. Both the difficulty in creating a saturated community and the more randomly dispersed trait patterns of randomly assembled communities reinforce the common trope in evolutionary literature that evolution an incredibly efficient process in organizing species in trait space.

Asymmetric competition can lead to both more interesting ecological (unstable or cyclic coexistence [Hernández-García *et al*., 2009]) and evolutionary dynamics in high dimensional phenotype space (e.g., Red-Queen dynamics [Rubin *et al*., 2021] or evolutionary chaos [Doebeli *et al*., 2017]), depending on the specific functional form of the competition kernel. While diversity patterns due to asymmetric competition is beyond the scope of this article, while investigating systems with non-stable (Red Queen) evolutionary dynamics we found that randomly assembling super-diverse communities was possible, but exceedingly rare [Rubin *et al*., 2021], creating a possible analogy to the difficulties in creating saturated communities with symmetric competition. We have also described a related phenomenon where continual selection can trap communities in low diversity meta-stable states [Rubin *et al*., 2021]. Taken together, these results suggest that if communities are unable to become saturated on ecological timescales during nascent community assembly (top-down assembly) or through successive invasions (bottom-up assembly) and evolution may, in certain cases, actually maintain unsaturated states rather than always pushing towards adaptive radiations, fully saturated ecological communities may be the exception rather than the rule.

## Acknowledgements

YI acknowledges support from FONDECYT (The National Fund for Scientific and Technological Development of Chile) project no. 1200708 (https://www.conicyt.cl/fondecyt/fondecyt-program/). MD acknowledges support from NSERC (The Natural Sciences and Engineering Research Council of Canada) grant no. 219930 (https://www.nserc-crsng.gc.ca/NSERC-CRSNG/Index_eng.asp)

## References

Allesina, S. & Tang, S. (2012). Stability criteria for complex ecosystems. Nature, 483, 205–208.

Allesina, S. & Tang, S. (2015). The stability-complexity relationship at age 40: a random matrix perspective. Population Ecology, 57, 63–75.

Baptestini, E.M., de Aguiar, M.A., Bolnick, D.I. & Araújo, M.S. (2009). The shape of the competition and carrying capacity kernels affects the likelihood of disruptive selection. Journal of Theoretical Biology, 259, 5–11.

Barabás, G. & D’Andrea, R. (2016). The effect of intraspecific variation and heritability on community pattern and robustness. Ecology Letters, 19, 977–986.

Barabás, G., Michalska-Smith, M.J. & Allesina, S. (2016). The effect of intra-and interspecific competition on coexistence in multispecies communities. The American Naturalist, 188, E1–E12.

Barbier, M., de Mazancourt, C., Loreau, M. & Bunin, G. (2021). Fingerprints of high-dimensional coexistence in complex ecosystems. Physical Review X, 11, 011009.

Chase, J.M. (2003). Community assembly: when should history matter? Oecologia, 136, 489–498.

Coyte, K.Z., Rao, C., Rakoff-Nahoum, S. & Foster, K.R. (2021). Ecological rules for the assembly of microbiome communities. PLOS Biology, 19, e3001116.

D’Andrea, R., Riolo, M. & Ostling, A.M. (2019). Generalizing clusters of similar species as a signature of coexistence under competition. PLOS Computational Biology, 15, e1006688.

Diaz-Colunga, J., Lu, N., Sanchez-Gorostiaga, A., Chang, C.Y., Cai, H.S., Goldford, J.E., Tikhonov, M. & Sánchez, Á. (2022). Top-down and bottom-up cohesiveness in microbial community coalescence. Proceedings of the National Academy of Sciences, 119, e2111261119.

Doebeli, M. (2011). Adaptive Diversification (MPB-48). Princeton University Press.

Doebeli, M. & Ispolatov, I. (2014). Chaos and unpredictability in evolution. Evolution, 68, 1365–1373.

Doebeli, M., Ispolatov, Y. & Simon, B. (2017). Towards a mechanistic foundation of evolutionary theory. eLife, 6, e23804.

Dougoud, M., Vinckenbosch, L., Rohr, R.P., Bersier, L.F. & Mazza, C. (2018). The feasibility of equilibria in large ecosystems: A primary but neglected concept in the complexity-stability debate. PLOS Computational Biology, 14, e1005988.

Débarre, F., Nuismer, S.L. & Doebeli, M. (2014). Multidimensional (co)evolutionary stability. The American Naturalist, 184, 158–171.

Edwards, K.F., Kremer, C.T., Miller, E.T., Osmond, M.M., Litchman, E. & Klausmeier, C.A. (2018). Evolutionarily stable communities: a framework for understanding the role of trait evolution in the maintenance of diversity. Ecology Letters, 21, 1853–1868.

Fowler, M.S. (2009). Increasing community size and connectance can increase stability in competitive communities. Journal of Theoretical Biology, 258, 179–188.

Fukami, T. & Nakajima, M. (2011). Community assembly: alternative stable states or alternative transient states? Ecology Letters, 14, 973–984.

Gilpin, M. (1994). Community-level competition: asymmetrical dominance. Proceedings of the National Academy of Sciences, 91, 3252–3254.

Hernández-García, E., López, C., Pigolotti, S. & Andersen, K.H. (2009). Species competition: coexistence, exclusion and clustering. Philosophical Transactions of the Royal Society A: Mathematical, Physical and Engineering Sciences, 367, 3183–3195.

Herron, M.D. & Doebeli, M. (2011). Adaptive diversification of a plastic trait in a predictably fluctuating environment. Journal of Theoretical Biology, 285, 58–68.

Hubbell, S.P. & Foster, R.B. (1986). Community Ecology, Harper and Row Publishers, chap. Biology, Chance, and History and the Structure of Tropical Rain Forest Tree Communities, pp. 314–329.

Ingram, T., Costa-Pereira, R. & Araújo, M.S. (2018). The dimensionality of individual niche variation. Ecology, 99, 536–549.

Kremer, C.T. & Klausmeier, C.A. (2017). Species packing in eco-evolutionary models of seasonally fluctuating environments. Ecology Letters, 20, 1158–1168.

Lawlor, L.R. (1980). Structure and stability in natural and randomly constructed competitive communities. The American Naturalist, 116, 394–408.

Lehman, C.L. & Tilman, D. (2000). Biodiversity, stability, and productivity in competitive communities. The American Naturalist, 156, 534–552.

Leimar, O., Sasaki, A., Doebeli, M. & Dieckmann, U. (2013). Limiting similarity, species packing, and the shape of competition kernels. Journal of Theoretical Biology, 339, 3–13.

Levine, J.M., Bascompte, J., Adler, P.B. & Allesina, S. (2017). Beyond pairwise mechanisms of species coexistence in complex communities. Nature, 546, 56–64.

MacArthur, R. (1970). Species packing and competitive equilibrium for many species. Theoretical Population Biology, 1, 1–11.

Macarthur, R. & Levins, R. (1967). The limiting similarity, convergence, and divergence of coexisting species. The American Naturalist, 101, 377–385.

MacArthur, R.H. (1965). Patterns of species diversity. Biological Reviews, 40, 510–533.

Mateo, R.G., Mokany, K. & Guisan, A. (2017). Biodiversity models: What if unsaturation is the rule? Trends in Ecology & Evolution, 32, 556–566.

May, R.M. (1972). Will a large complex system be stable? Nature, 238, 413–414.

Meszéna, G., Gyllenberg, M., Pásztor, L. & Metz, J.A. (2006). Competitive exclusion and limiting similarity: A unified theory. Theoretical Population Biology, 69, 68–87.

Olszewski, T.D. (2011). Persistence of high diversity in non-equilibrium ecological communities: implications for modern and fossil ecosystems. Proceedings of the Royal Society B: Biological Sciences, 279, 230–236.

Pacala, S.W. & Roughgarden, J. (1982). The evolution of resource partitioning in a multidimensional resource space. Theoretical Population Biology, 22, 127–145.

Rael, R.C., D’Andrea, R., Barabás, G. & Ostling, A. (2018). Emergent niche structuring leads to increased differences from neutrality in species abundance distributions. Ecology, 99, 1633–1643.

Roughgarden, J. (1976). Resource partitioning among competing species—a coevolutionary approach. Theoretical Population Biology, 9, 388–424.

Rubin, I.N., Ispolatov, I. & Doebeli, M. (2021). Evolution to alternative levels of stable diversity leaves areas of niche space unexplored. PLOS Computational Biology, 17, e1008650.

Sasaki, A. & Ellner, S. (1995). The evolutionarily stable phenotype distribution in a random environment. Evolution, 49, 337.

Scheffer, M. & van Nes, E.H. (2006). Self-organized similarity, the evolutionary emergence of groups of similar species. Proceedings of the National Academy of Sciences, 103, 6230–6235.

Schluter, D. (2000). The Ecology of Adaptive Radiation. Oxford University Press.

Serván, C.A. & Allesina, S. (2021). Tractable models of ecological assembly. Ecology Letters, 24, 1029–1037.

Serván, C.A., Capitán, J.A., Grilli, J., Morrison, K.E. & Allesina, S. (2018). Coexistence of many species in random ecosystems. Nature Ecology & Evolution, 2, 1237–1242.

Sigmuiud, K. (1995). Darwin’s “circles of complexity”: Assembling ecological communities. Complexity, 1, 40–44.

Song, C. & Saavedra, S. (2018). Will a small randomly assembled community be feasible and stable? Ecology, 99, 743–751.

Svanbäck, R., Pineda-Krch, M. & Doebeli, M. (2009). Fluctuating population dynamics promotes the evolution of phenotypic plasticity. The American Naturalist, 174, 176–189.

Szabó, P. & Meszéna, G. (2006). Limiting similarity revisited. Oikos, 112, 612–619.

