## Supplementary Information for "Ecological diversity exceeds evolutionary diversity in model ecosystems"

March 24, 2022

#### A Feasibility of random species

After randomly assembling a founding population we calculate the probability this initial random assortment of species is feasible. As we only consider Gaussian competition, any feasible community will also be globally stable [Hernández-García *et al.*, 2009]. The probability $N$ randomly selected, competitive species will be feasible always decreases with $N$. Additionally, the probability those $N$ species are feasible also always increases with the dimension of competition as the available niche space for those $N$ species is also greater.

When the probability of feasibility curves for each dimension are re-scaled to be a function of the community saturation ($H/ESS^d$) instead of diversity ($H$), the probability curves take on a remarkably similar shape regardless of dimension when viewed on a log-log scale [Figure C6] and it become clear that while the probability of feasibility for $N$ random species increases with dimension, the probability of feasibility for a random community of a given saturation actually decreases with increasing dimension. Notably, for $\sigma_\alpha = 0.6$ only in dimensions 1 and 2 were we able to randomly assemble a saturated, feasible community. In dimension 1, 34.77% of all 3 species communities (i.e., saturated) were feasible, and 9.54%, 0.44%, and 0.02% of communities with 4, 5, and 6 species (super-saturated) were found to be feasible respectively. In dimension 2, only 0.50% of 9 species communities (i.e., fully saturated) were feasible, and 0.08%, 0.06%, and 0.01% of 10-12 species (super-saturated) communities respectively were found to be feasible. While it is certainly possible to randomly select $N$ species that represent both a saturated or super-saturated and feasible community, in higher dimensions the probability of doing so is essentially zero [Figure C7].

##### Universal scaling of feasibility probability

The curves in Figures C6 and C8 corresponding to various dimensions looks similar to each other. Indeed, it turns out that for every $\sigma_\alpha$, and thus every unique ESS, there exists a scaling across dimensions that makes distributions of feasibility universal: The probability distribution of a randomly assembled community of a given saturation compared to the ESS being feasible scales as $a^{-d}$ [Figure C9]. The constant $a$ is non-universal and depends on $\sigma_\alpha$. This is because for each $\sigma_\alpha$ and thus for each ESS, there is a certain set of domains in phenotype space where the species must be located for the community to be viable [Figure 1]. The random assembly of species results in a given probability to have a viable community of a certain number of species for a given set of domains, that is, for a given $\sigma_\alpha$. The fraction of total phenotype space with dimension $d$ that corresponds to these domains scales as $a^{-d}$, which explains the observed scaling. Yet for a new $\sigma_\alpha$, such a set of domains changes in non-monotonic way due to the emergence of new species and resulting appearance, increase or shrinking of certain domains. Hence, the scaling factor is non-universal (in $\sigma_\alpha$) and even does not depend on $\sigma_\alpha$ in any systematic way.

### B Results of Statistical Test for Trait Dispersion in One Dimension

To test whether stable communities emerging through random assembly we used Welch's t-test to compare top-down and bottom-up communities to Gaussian and uniformly distributed species of the same diversity as the communities. We also compared top-down and bottom-up communities to each other. We are using the coefficient of variation of the distance to the nearest neighbor in trait space ($CV_{NN}$) for each species as our metric for trait dispersion. Perfectly ordered communities (those that form a lattice in trait space) are maximally under dispersed compared to random and will have a $CV_{NN} = 0$. As communities become more dispersed in trait space they will have a larger $CV_{NN}$. As a metric of dispersion is meaningless in communities with 2 species, communities with diversity 3 through 6 (the maximal diversity for feasible communities with 1-dimensional competition and a $\sigma_\alpha = 0.6$) were tested.

Communities built through top-down and bottom-up assembly processes were not found to differ the the dispersion of their traits [Table B1]. However, both types of randomly assembled community were found to be significantly ($\alpha < 0.05$) more regularly spaced in trait space compared to both methods of randomly distributed traits for all levels of diversity.

Because of computational limitations we did not statistically test the trait dispersion for communities with competition in greater than one dimension. However, graphically these communities have noticeably lower values $CV_{NN}$ compared both to both forms of randomly distributed traits of the same dimension, alligning with the statistical results from dimension one.

| **Diversity** | **Model 1** | **Model 2** | **p value** | **Model 1 mean $CV_{NN}$** | **Model 2 mean $CV_{NN}$** |
|---|---|---|---|---|---|
| 3 | Top Down | Bottom Up | 0.178 | 0.337 | 0.191 |
| 4 | Top Down | Bottom Up | 0.866 | 0.443 | 0.432 |
| 5 | Top Down | Bottom Up | 0.253 | 0.614 | 0.560 |
| 6 | Top Down | Bottom Up | 0.427 | 0.572 | 0.621 |
| 3 | Top Down | Gaussian Distributed | $3.14 \times 10^{-2}$ * | 0.337 | 0.646 |
| 4 | Top Down | Gaussian Distributed | $7.96 \times 10^{-5}$ ** | 0.443 | 0.814 |
| 5 | Top Down | Gaussian Distributed | $1.88 \times 10^{-2}$ * | 0.614 | 0.823 |
| 6 | Top Down | Gaussian Distributed | $1.68 \times 10^{-4}$ ** | 0.572 | 0.923 |
| 3 | Bottom Up | Gaussian Distributed | $7.09 \times 10^{-4}$ ** | 0.191 | 0.646 |
| 4 | Bottom Up | Gaussian Distributed | $5.25 \times 10^{-5}$ ** | 0.432 | 0.814 |
| 5 | Bottom Up | Gaussian Distributed | $3.96 \times 10^{-3}$ ** | 0.560 | 0.823 |
| 6 | Bottom Up | Gaussian Distributed | $8.26 \times 10^{-4}$ ** | 0.621 | 0.923 |
| 3 | Top Down | Uniformly Distributed | $9.66 \times 10^{-3}$ ** | 0.337 | 0.700 |
| 4 | Top Down | Uniformly Distributed | $3.87 \times 10^{-3}$ ** | 0.443 | 0.696 |
| 5 | Top Down | Uniformly Distributed | $3.88 \times 10^{-2}$ * | 0.614 | 0.782 |
| 6 | Top Down | Uniformly Distributed | $3.20 \times 10^{-4}$ ** | 0.572 | 1.04 |
| 3 | Bottom Up | Uniformly Distributed | $8.81 \times 10^{-5}$ ** | 0.191 | 0.700 |
| 4 | Bottom Up | Uniformly Distributed | $2.67 \times 10^{-3}$ ** | 0.432 | 0.696 |
| 5 | Bottom Up | Uniformly Distributed | $7.05 \times 10^{-3}$ ** | 0.560 | 0.782 |
| 6 | Bottom Up | Uniformly Distributed | $9.11 \times 10^{-4}$ ** | 0.621 | 1.04 |

**Table B1: Statistical Test for Trait Dispersion in One Dimension** To test for the dispersion of traits, we calculated the coefficient of variation of the nearest neighbor for each species in the communities ($CV_{NN}$). Communities randomly assembled through both top-down and bottom-up assembly processes were tested against random uniformly and Gaussian distributed traits as well as against each other using Welch's t-test. One asterisks denotes significance with $\alpha < 0.05$ and two denotes significance for $\alpha < 0.01$. Please see Figure 6 for a graphical representation of the data tested.

### C Supplementary figures

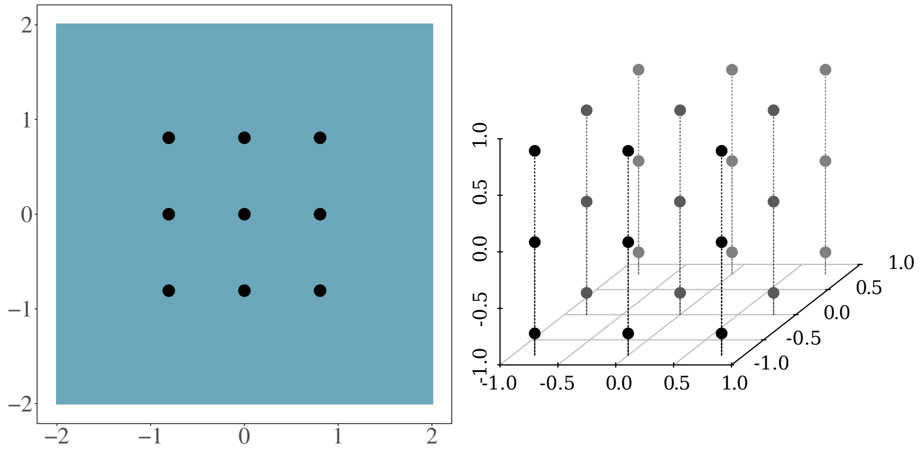


**(a)** 2-dimensional ESS and invasion fitness **(b)** 3-dimensional ESS

**Figure C1: ESS in dimension two and three.** Evolutionary stable states were calculated using adaptive dynamics. Panel A shows the ESS in 2D. Blue represents areas with negative invasion fitness. As the ESS is globally stable, no areas of positive invasion fitness exist. Panel B the ESS in 3D. $\sigma_\alpha = 0.6$

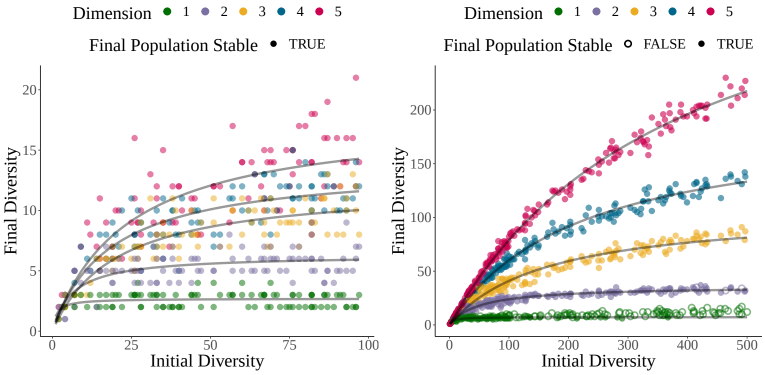


**(a)** $\sigma_\alpha = 1$ $(H_{ESS} = 2^d)$ **(b)** $\sigma_\alpha = 0.5$ $(H_{ESS} = 4^d)$

**Figure C2: The final stable diversity of a randomly assembled community for $\sigma_\alpha = 0.5, 1.0$.** This is the same simulations as in Figure 3 but for $\sigma_\alpha = 1$ and 0.5 for panels A and B respectively. Communities were assembled with random species in 1-5 dimensions. Ecological dynamics were run for $10^8$ time steps. Hollow points represent communities that did not stabilize to a feasible community by the end of the simulation and still contain at least one species that will eventually go extinct. Grey lines are a regression using a nonlinear least squares fit to Michaelis-Menten dynamics ($y = \frac{ax}{b+x}$). Regressions were fit only to communities that have stabilized by the end of the simulation. The Michaelis-Menten curves are not mechanistic, and where not chosen to model the ecological dynamics in any way, but are asymptotic curves (asymptote equal to regression parameter $a$) that are a good visual fit for the data.

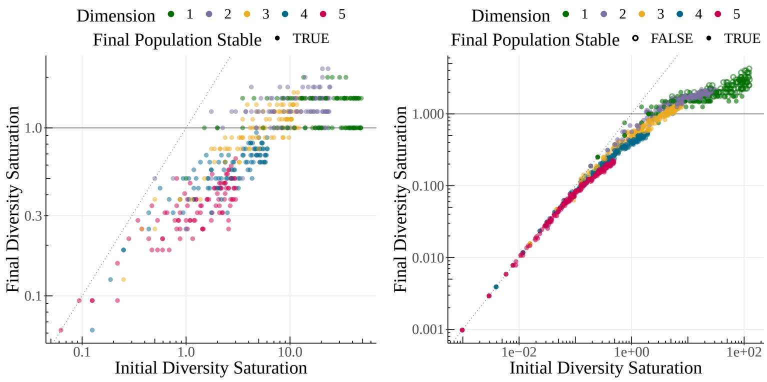


**(a)** $\sigma_\alpha = 1$ ($H_{ESS} = 2^d$) **(b)** $\sigma_\alpha = 0.5$ ($H_{ESS} = 4^d$)

**Figure C3: Final community saturation from randomly assembled species for $\sigma_\alpha = 0.5, 1.0$.** This is the same simulations as in Figure 3 but for $\sigma_\alpha = 1$ and 0.5 for panels A and B respectively. Communities were assembled with random species in 1-5 dimensions. Ecological dynamics were run for $10 \times 10^8$ time steps. Community saturation is measured as the diversity divided by the diversity of the ESS for the same ecosystem (for $\sigma_\alpha = 0.5$, $H_{ESS} = 4^d$, for $\sigma_\alpha = 1$, $H_{ESS} = 2^d$).

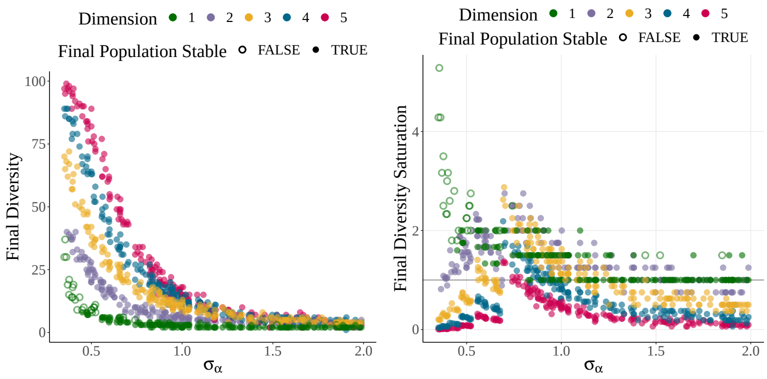


**(a)** Community diversity **(b)** Community saturation

**Figure C4: Community diversity and saturation as a function of the width of competition ($\sigma_\alpha$).** Random communities of 100 species were assembled and ecological dynamics were run for $10^{12}$ time steps. As expected, as the width of competition ($\sigma_\alpha$) increases (species are more generalist), the final diversity of the community decreases. However, the saturation of the communities peaks at an intermediate value of $\sigma_\alpha$. This is because two communities with similar $\sigma_\alpha$ will likely share the same diversity of their ESS, but the community with a larger $\sigma_\alpha$ will have a smaller community on average when that community is assembled randomly. Colors represent communities competing in different numbers of dimensions. Hollow points represent communities that did not stabilize when the simulation concluded. Quartic carrying capacity function ($\sigma_K = 1$) and Gaussian competition kernel ($\sigma_\alpha = 0.6$)

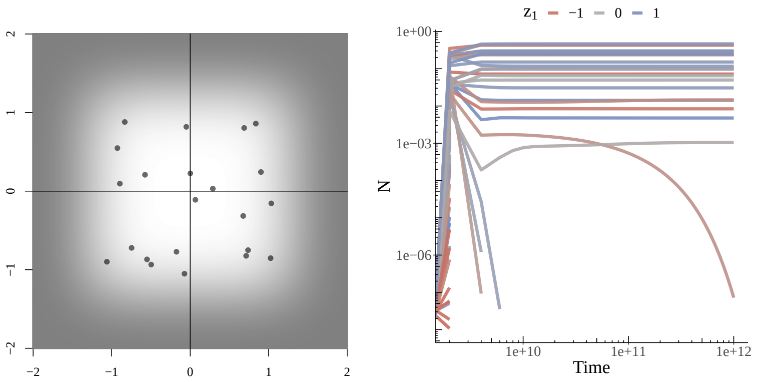


**(a)** Final community

**(b)** Population size of each species in the community

**Figure C5: Example of community with transient dynamics.** An example of a community with transient dynamics. After $10^{12}$ time steps the community had not yet stabilized. Founding population contained 311 species with random phenotypes in 2-dimensional phenotype space. Panel A shows the final population configuration and the carrying capacity function (white = 1, darker means decreasing $K$). Panel B shows the population size of each species over the course of the simulation. The species that is undergoing transient extinction at the end of the simulation has a phenotype of $[-0.496, -0.935]$. Quartic carrying capacity and Gaussian competition function with $\sigma_\alpha - 0.6$.

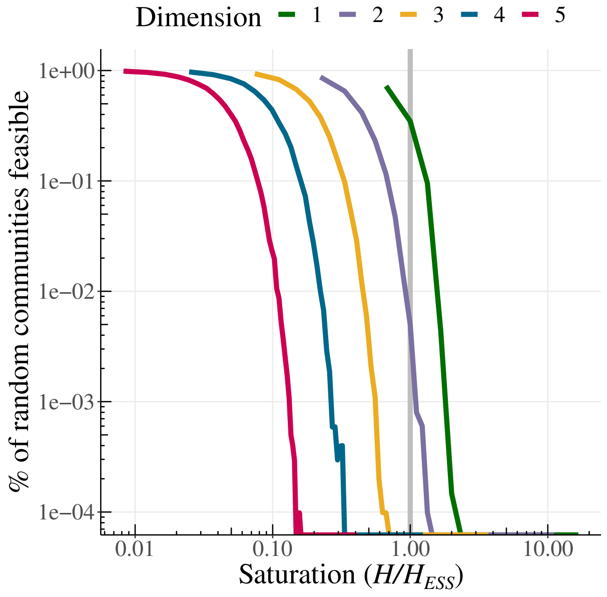


**Figure C6: Probability randomly assembled communities of a given saturation are feasible.** Each color represents the dimension of the trait space (from $1-5$). The grey line denotes the saturation point where the randomly assembled community has the same diversity as the ESS in that dimension. Each probability curve was generated by calculating the feasibility of $10^6$ communities with between 2 and 100 randomly chosen species for each dimension (2 to 50 for dimension 1). On average 10200 communities were simulated for each level of diversity in each dimension (double that for dimension 1), allowing us to calculate probabilities to the order of $10^{-4}$. Quartic carrying capacity function ($\sigma_K = 1$) and Gaussian competition kernel with $\sigma_\alpha = 0.6$.

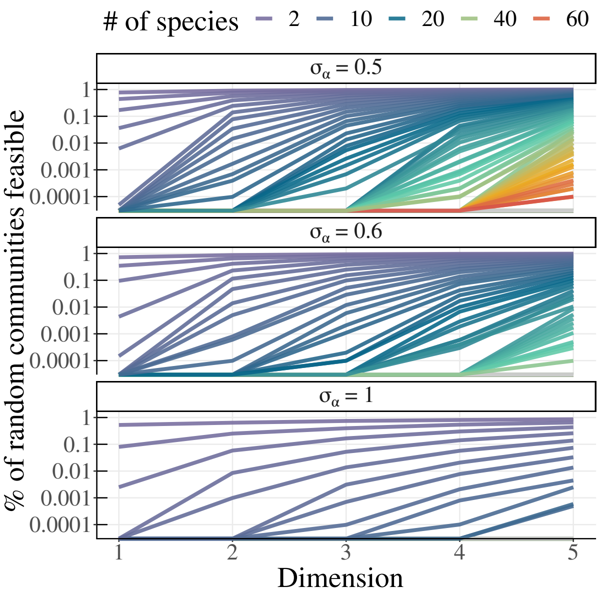


**Figure C7: Probability that randomly assembled communities of a given diversity are feasible as a function of the dimension.** Each line (and color) represents communities of a given diversity. Curves were generated by calculating the feasibility of $\times 10^6$ communities with between 2 and 100 randomly chosen species (between 2 and 50 for 1D) for each dimension. On average 10200 communities (twice that for 1D) were simulated for each level of diversity in each dimension, allowing us to calculate probabilities to the order of $1 \times 10^{-4}$. Quartic carrying capacity function ($\sigma_K = 1$) and Gaussian competition kernel ($\sigma_\alpha = 0.6$)

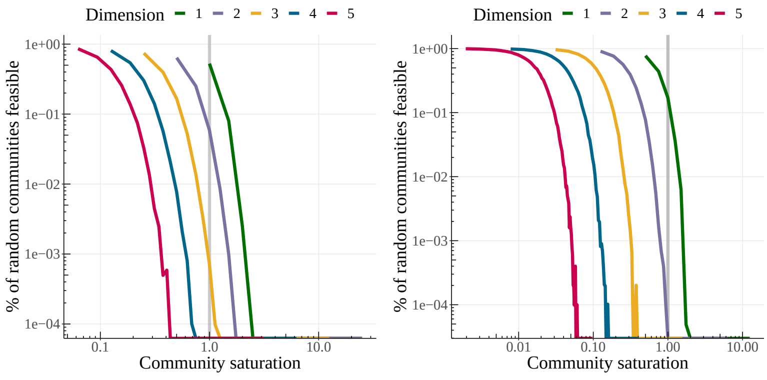


**(a)** $\sigma_\alpha = 1$ ($H_{ESS} = 2^d$) **(b)** $\sigma_\alpha = 0.5$ ($H_{ESS} = 4^d$)

**Figure C8: Probability randomly assembled communities of a given saturation compared to the ESS are feasible.** This is the same simulations as in Figure C6 but for $\sigma_\alpha = 1$ and 0.5 for panels A and B respectively. Each color represents the dimension of the trait space (from $1-5$). The grey line denotes the saturation point where the randomly assembled community has the same diversity as the ESS in that dimension. Each probability curve was generated by calculating the feasibility of $10^6$ communities with between 2 and 100 randomly chosen species for each dimension (2 to 50 for dimension 1). On average 10200 communities were simulated for each level of diversity in each dimension (double that for dimension 1), allowing us to calculate probabilities to the order of $10^{-4}$.

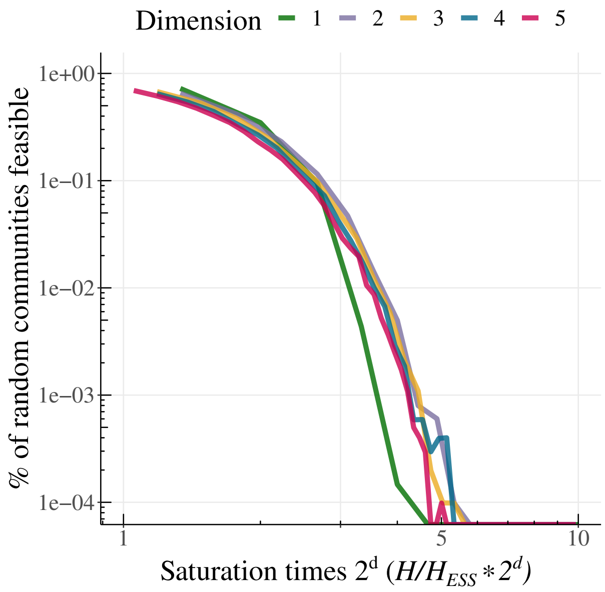


**Figure C9: Universal scaling of probability that randomly assembled communities of a given saturation are feasible.** Each color represents the dimension of the trait space (from $1-5$). The saturation of each community is multiplied by $2^d$ to illustrate the approximate universality of the curves. Just as in Figure 4, each probability curve was generated by calculating the feasibility of $10^6$ communities with between 2 and 100 randomly chosen species for each dimension (2 to 50 for dimension 1). On average 10200 communities were simulated for each level of diversity in each dimension (double that for dimension 1), allowing us to calculate probabilities to the order of $10^{-4}$. Quartic carrying capacity function ($\sigma_K = 1$) and Gaussian competition kernel with $\sigma_\alpha = 0.6$ were used.
